# Junction-level imaging and modeling of transient leaks in epithelia predicts a dynamic tight junction strand network

**DOI:** 10.64898/2026.11.06.764832

**Authors:** Anni Pörsti, Aapo Tervonen, Huy Tran, Soile Nymark, Teemu O. Ihalainen

## Abstract

Epithelial tight junctions restrict paracellular permeability, but how small solutes leak through the barrier is not fully understood, because permeability is usually measured globally. Here, we combined fast zinc-based barrier imaging (ZnUMBA) with computational modelling of the dynamic tight junction strand network to study transient leaks at single bicellular junctions in MDCK II cells. Leaks were short-lived, with a median duration of 17 s in wild-type cells, and occurred in all cell lines studied, including ZO-1 knockout, ZO-2 knockout, and ZO-1-overexpressing cells. In ZO-2 knockout and ZO-1-overexpressing cells, leaks lasted longer without changes in their frequency or transepithelial electrical resistance (TER). ZO-1 knockout increased the junctional ZnUMBA signal and reduced TER and the number of detected leaks. Simulations showed that stochastic breaking and sealing of tight junction strands is sufficient to explain the observed leak frequencies and durations, and that a single pair of strand break formation and sealing rates reproduces both junction-level leaks and global permeability. Whereas leak frequency mainly reflects the ratio of these rates, leak duration provides independent information on strand sealing that global measurements cannot resolve. Modulating myosin II activity had little effect on leaks, whereas actin depolymerization compromised the barrier. Together, these results link junction-level leaks of ions and small solutes to tight junction strand dynamics.

## Introduction

Epithelia line body surfaces and organs, forming barriers that separate body compartments. These barriers are formed by tight junctions, located near the apical side of the lateral membrane between neighboring cells. Tight junctions are large protein complexes where transmembrane proteins such as claudins and occludin form two-dimensional, anastomosing strand networks on the cell membrane, pairing with strands of the adjacent cell (Fuladi et al., 2022; Otani et al., 2019; Samanta et al., 2018; Staehelin, 1973). Claudins are connected to the cytoplasmic zonula occludens proteins ZO-1 and ZO-2, which function as scaffolds and signaling hubs (Zihni et al., 2016). ZO-1 is a major scaffolding protein found in epithelia (Beutel et al., 2019), and it connects tight junctions to the actin cytoskeleton via its C-terminal actin-binding region (Itoh et al., 1997; Schwayer et al., 2019; Spadaro et al., 2017; Van Itallie et al., 2017). This connection enables its C-terminal region to unfold in a mechanosensitive manner (Spadaro et al., 2017), and the affinity of actin binding has been shown to affect the epithelial barrier (Belardi et al., 2020). However, the role of ZO proteins and the actomyosin cytoskeleton in tight junction barrier regulation remains incompletely understood.

Tight junctions control the paracellular permeability of ions and molecules via two pathways. First, ions and small solutes diffuse via the so-called pore pathway formed by different claudins and regulated by their expression (Günzel and Yu, 2013; Tsukita et al., 2019; Zihni et al., 2016). Second, the so-called leak pathway is a low-capacity route that allows the passage of larger molecules in addition to ions and small solutes (Anderson and Van Itallie, 2009; Monaco et al., 2021; Shen et al., 2011; Zihni et al., 2016). The precise origin of the leak pathway, however, is not fully known. Central tubes formed in the tricellular junctions have been suggested to allow macromolecules to cross the epithelium, but their contribution to the overall epithelial permeability is relatively low due to their low density (Krug et al., 2009). For bicellular junctions, the so-called dynamic strand model proposes that strands constantly break and seal, allowing slow, stepwise diffusion through the network (Otani and Furuse, 2020; Tervonen et al., 2019). Strand networks are indeed highly dynamic in claudin-expressing fibroblasts (Sasaki et al., 2003; Van Itallie et al., 2017). Occasionally, breaks in successive strands may also coincide, transiently opening a path through the entire, otherwise intact network (Tervonen et al., 2019). In contrast, Stephenson et al. (2019) observed transient, junction-scale barrier openings in the mechanically active epithelium of *Xenopus laevis* embryos, where tension appears to disrupt the junction and the breach is actively repaired by RhoA activation and actomyosin contraction, termed Rho flares.

Usually, epithelial permeability is measured with global methods such as transepithelial electrical resistance (TER) or dextran permeability assays. These methods average over the whole monolayer and over time, and therefore cannot resolve barrier dynamics at the level of individual junctions. Recently, methods have been established to provide a more local, junction-level view of the barrier and its dynamics (Higashi et al., 2023; Richter et al., 2022; Stephenson et al., 2019). Stephenson et al. (2019) used the zinc-based ultrasensitive microscopic barrier assay (ZnUMBA) to detect junction-scale barrier openings in *Xenopus* embryos. In ZnUMBA, zinc ions (Zn^2+^) and a cell-impermeable zinc indicator are placed on opposite sides of the epithelium. The indicator fluoresces only when bound to Zn^2+^, so transient local increases in junctional signal, termed leaks, reveal where and when Zn^2+^ crosses the barrier at a single bicellular junction (Higashi et al., 2023; Stephenson et al., 2019). ZnUMBA has since been applied to Madin-Darby canine kidney (MDCK) II cells, where leaks were also detected, and their repair was shown to involve proteolytic cleavage of EpCAM and release of claudin-7 (Higashi et al., 2022). Importantly, tension-induced breaches and stochastic open paths through the strand network would both appear in ZnUMBA as a similar transient increase in junctional signal. Which of these mechanisms underlies the leaks observed in MDCK cells, what determines their frequency and duration, and how they relate to global barrier properties therefore remain unknown.

As scaffolds that organize claudins into strands (Umeda et al., 2006), ZO proteins are natural candidates for regulating strand dynamics and junction-level leaks. The roles of the ZO proteins in barrier maintenance have been studied using knockdown (KD) and knockout (KO) cell lines. In MDCK cells, KO of either ZO-1 or ZO-2 caused no obvious barrier defects, apart from increased permeability for 4 kDa dextran in ZO-1 KO cells (Otani et al., 2019; Tokuda et al., 2014). Similarly, ZO-1 KD in MDCK cells selectively impaired the barrier to large solutes without affecting small solute flux through the claudin pores and altered the TER response to myosin inhibition (Van Itallie et al., 2009). In contrast, ZO-1/ZO-2 double KO cells lacked tight junction strands, showed markedly decreased TER and increased paracellular flux regardless of molecular size, and had disorganized epithelial polarity (Otani et al., 2019). Thus, ZO-1 and ZO-2 are partially redundant, and at least one of them is required for tight junction formation (Otani et al., 2019; Umeda et al., 2006). However, the barrier was assessed with global measurements, and how ZO proteins and their link to the actomyosin cytoskeleton affect the frequency and duration of junction-level leaks is unknown.

In this study we used ZnUMBA to measure junction-level permeability of claudin-2-expressing MDCK II cells. We compared wild type (WT), ZO-1 KO, ZO-2 KO, and ZO-1 overexpression (OE) cell lines and found leaks in all of them. Observed leak frequency was lower in ZO-1 KO cells, whereas leaks lasted longer in ZO-2 KO and ZO-1 OE cells. Using a computational model of the dynamic strand network, we show that the observed leak frequencies and durations are consistent with stochastic breaking and sealing of tight junction strands, and predict that ZO-2 KO and ZO-1 OE shift strand dynamics toward fewer but longer-lasting breaks without changing global barrier properties. Pharmacological modulation of myosin II activity had little effect on leak behavior in WT and ZO-1 KO cells, whereas actin depolymerization compromised the barrier and reduced detected leak frequency in WT cells.

## Results

### Characterization of ZO protein cell lines and the ZnUMBA setup

To study how ZO proteins affect junction-level barrier dynamics, we used MDCK II WT cells together with lines lacking ZO-1 or ZO-2 (ZO-1 KO and ZO-2 KO), and a line expressing ZO-1 fused to mRuby in the WT background. We originally established the ZO-1 mRuby line to visualize tight junctions during ZnUMBA imaging. Because the fusion protein is expressed in addition to endogenous ZO-1, and flow cytometry showed increased total ZO-1 levels (1.23-fold, SFig. 1A), we used it as a ZO-1 OE line.

Immunostaining of tight junction proteins confirmed the genotypes of the cell lines: ZO-1 KO cells lacked ZO-1 staining, and ZO-2 KO cells lacked junctional ZO-2 staining (Fig. 1C). Claudin-2, a major claudin in MDCK II cells, localized to cell junctions in all lines, but its junctional localization was noticeably reduced in ZO-1 KO cells, consistent with previous observations (Haas et al., 2022). ZO-2 staining was weak and largely cytoplasmic in all lines, but a faint junctional signal was detected in WT, ZO-1 KO and ZO-1 OE cells. In WT and ZO-1 OE cells, claudin-2 and ZO-1 colocalized with junctional actin. Thus, junctional claudin-2 and ZO-2 indicated that tight junctions still formed in ZO-1 KO cells, consistent with either ZO protein being sufficient for tight junction formation (Otani et al., 2019; Umeda et al., 2006), although loss of ZO-1 reduced junctional claudin-2.

**Figure 1.**
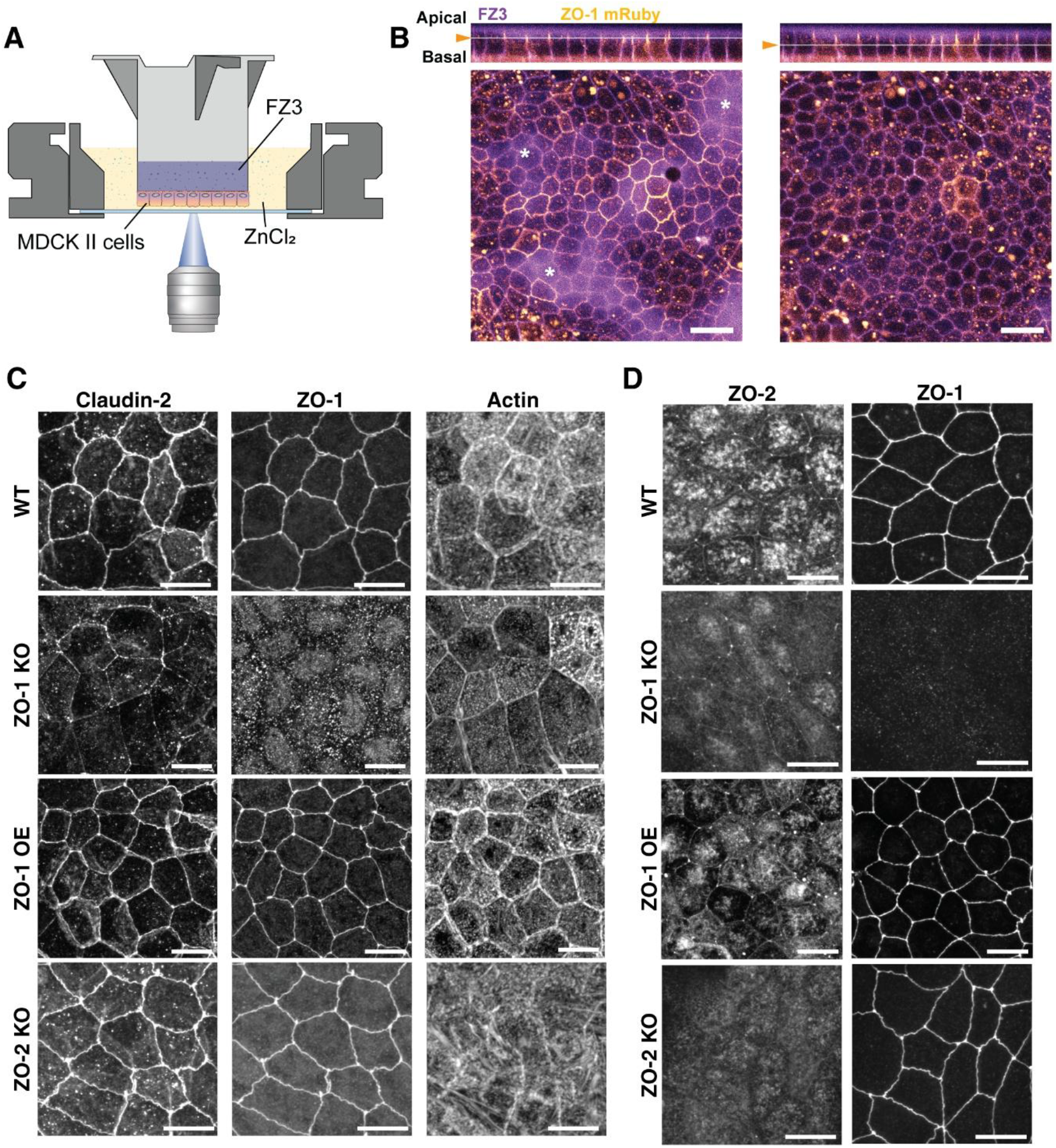
ZnUMBA setup and characterization of the cell lines. (A) Schematic of the ZnUMBA setup. MDCK II cells were grown on the underside of a filter insert. FluoZin-3 (FZ3) was added to the basal compartment inside the insert and ZnCl2 to the apical compartment in the well, and the cells were imaged from below with an inverted confocal microscope. (B) ZnUMBA (FZ3, magenta) and ZO-1 mRuby (yellow) signals in ZO-1 OE cells, shown as a top view from two depths with orthogonal x-z sections. ZO-1 mRuby localizes near the apical end of the lateral membrane. Basal to it, the ZnUMBA signal is confined to the lateral intercellular space and outlines the cell borders (right), whereas apical to it the signal is diffuse (left, white asterisks). Orange arrow and white line in both x-z section indicate x-y image plane. (C) Immunostaining of claudin-2, ZO-1 and F-actin (phalloidin) in WT, ZO-1 KO, ZO-1 OE and ZO-2 KO cells. (D) ZO-2 and ZO-1 immunostaining shown as projection from apical slices from WT, ZO-1 KO, ZO-1 OE and ZO-2 KO cells. Scale bars 20 µm (B) and 10µm (C and D).

To measure barrier dynamics at individual junctions, we performed ZnUMBA on MDCK monolayers grown on the underside of filter inserts (Fig. 1A) (Higashi et al., 2023; Stephenson et al., 2019). In this inverted configuration, ZnCl_2_ was added to the apical compartment in the well and the zinc indicator FluoZin-3 (FZ3) to the basal compartment inside the insert. To locate the tight junctions relative to the ZnUMBA signal, we located ZO-1 mRuby expression from x-z sections of 3D image stacks. ZO-1 mRuby localized near the apical end of the lateral membrane (Fig. 1B). Basal to the tight junction, the ZnUMBA signal was confined to the narrow lateral intercellular space and outlined the cell borders. Apical to the tight junction, the signal spread diffusely, as indicator that had reached the Zn^2+^-containing apical medium was no longer confined between cells. Because cell height varied,we set the imaging plane for each field just basal to the tight junctions, where the junctional signal was confined and individual bicellular junctions could be resolved. In this plane, the ZnUMBA signal outlined the same cell borders as ZO-1 mRuby (Fig. 1B).

### Transient leaks occur at bicellular junctions in all cell lines

We recorded 10-min time-lapse videos of the ZnUMBA signal in all cell lines with identical imaging settings and the same focal-plane criterion. Because the junctional ZnUMBA signal itself outlined the cell borders, ZO-1 mRuby was not needed to locate the junctions, which allowed all lines to be analyzed in the same way. We segmented the cell borders from the ZnUMBA signal using Cellpose (Stringer et al., 2021) and measured the mean ZnUMBA intensity along the segmented bicellular junctions over the whole recording, including any leaks. The mean absolute junctional intensity was lowest in WT cells, approximately 2-fold higher in ZO-2 KO and ZO-1 OE cells, and approximately 4-fold higher in ZO-1 KO cells (SFig. 1B; p < 0.0001 for all comparisons with WT).

As previously reported for MDCK II cells (Higashi et al., 2023, 2022), WT monolayers showed leaks: transient increases in junctional ZnUMBA intensity that returned to baseline (Fig. 2A′ and A″). To detect leaks, we followed the intensity of each segmented junction over time. A leak was defined as an increase in junctional intensity of at least 20% above the local baseline (the lowest intensity within a surrounding 50-s window), lasting at least 5 s. Because the threshold is relative to the baseline, leaks of the same absolute size are harder to detect when the junctional signal is high, as in ZO-1 KO cells (Fig. 2E′ and E″).

**Figure 2.**
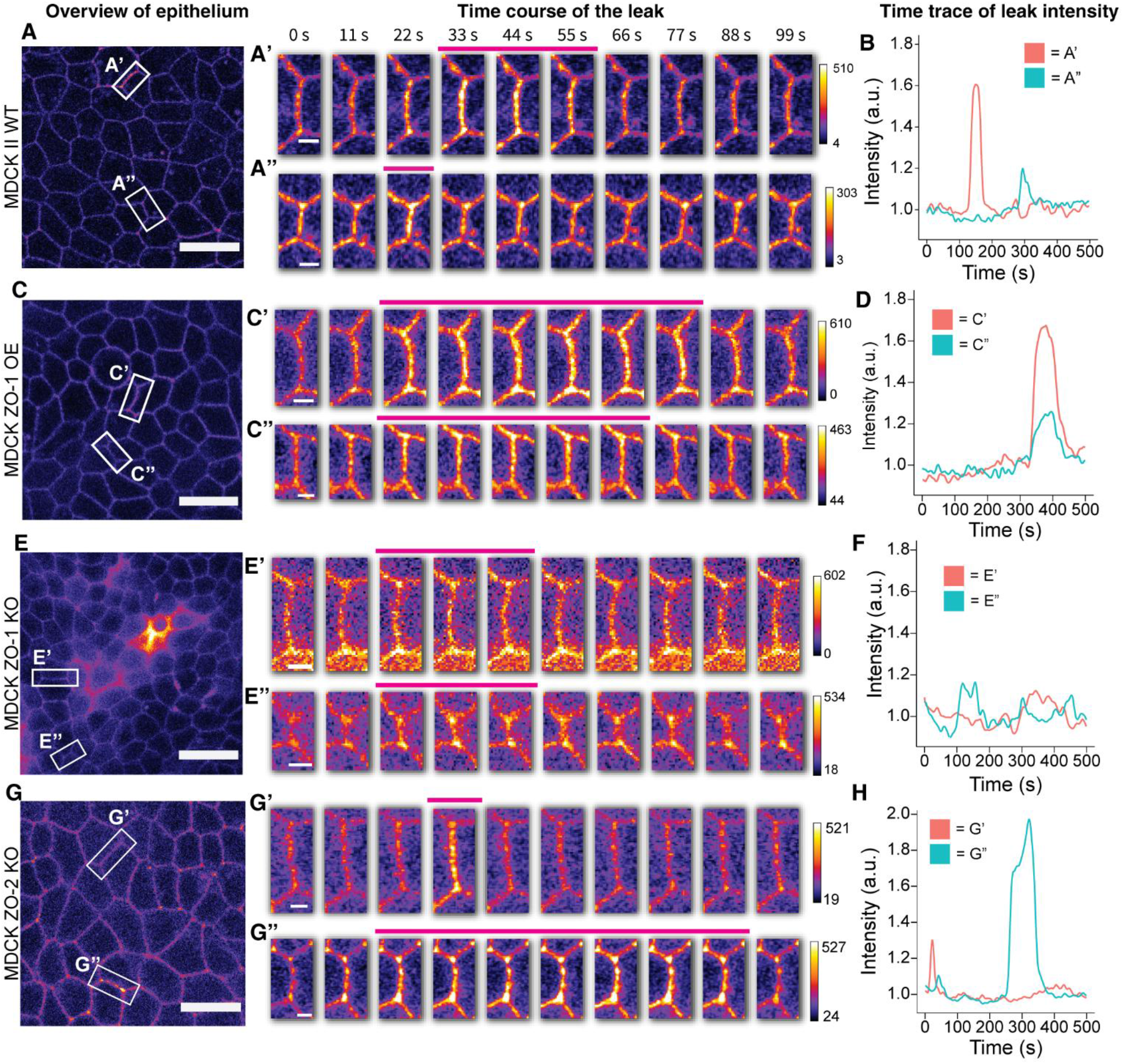
Transient leaks at bicellular junctions in MDCK II cell lines. (A, C, E, and G) Overview of the ZnUMBA (FZ3) signal in WT (A), ZO-1 OE (C), ZO-1 KO (E) and ZO-2 KO (G) monolayers. The junctional signal outlines the cell borders. White boxes mark the junctions shown in the time courses. (A′, A″, C′, C″, E′, E″, G′, and G″) Time courses of representative leaks at the boxed junctions, shown every 11 s (every tenth frame) for respective cell lines. Pink bars indicate the frames in which the junctional intensity exceeded the leak threshold. (B, D, F, and H) Normalized junctional intensity of the junctions in the corresponding time courses over 500 s of the recording. Intensities were normalized to the mean junctional intensity excluding values above the leak threshold. Scale bars, 20 µm (A, C, E, and G) and 3 µm (B, D, F, and H).

Leaks occurred in all cell lines (Fig. 2A-H). In WT, ZO-1 OE, and ZO-2 KO cells, leaks often appeared as distinct transient increases in junctional intensity, whereas in ZO-1 KO cells they were less distinct against the higher junctional signal (Fig. 2E and F, SFig. 1B). Analysis of junction subsections showed that during most leaks, the intensity increased along the whole junction rather than being concentrated toward one end (data not shown). Leaks varied in duration and intensity within each cell line (Fig. 2B, D, F, and H). Thus, leaks were a common feature of MDCK II monolayers regardless of ZO protein levels. We did not observe junctional elongations preceding the leaks (SVideo 1 and 2).

In addition to transient leaks, tricellular junctions often showed a persistently higher ZnUMBA signal than the adjacent bicellular junctions. To quantify this, we calculated the ratio of the mean intensity at each tricellular junction to that of its neighboring bicellular junctions that showed no leaks during the video. A mean relative intensity was calculated for each video for analysis. The ratio was above 1 in all lines, highest in WT cells (1.472 ± 0.161, mean ± SD) and lower in ZO-1 OE (1.407 ± 0.078), ZO-2 KO (1.363 ± 0.092), and ZO-1 KO cells (1.254 ± 0.107) (SFig. 1C).

### Leak duration, frequency and TER in cell lines with altered ZO protein levels

To compare leak behavior between the cell lines, we quantified the duration and frequency of leaks in each time-lapse. We measured leak duration as the time the junctional intensity remained above the threshold. Only leaks that both started and ended within the recording were included in the duration analysis. Leaks still ongoing at the end of the recording (19.6–37.3% of leaks, depending on the cell line, STable 1) were excluded from the duration analysis. Because the number of detected leaks depends on the recording time and on the number of junctions in the field of view, we expressed leak frequency as the number of leaks per 100 bicellular junctions per minute.

Most leaks were short. In WT cells, leak durations ranged from 6.37 s to 342 s, with a median of 17.0 s (Fig. 3A and A′). Leaks in ZO-1 KO cells had a similar median duration (15.4 s), whereas leaks lasted longer in ZO-2 KO (24.4 s) and ZO-1 OE cells (32.9 s; both p < 0.0001 compared with WT), reflected in broader duration distributions shifted toward longer values (Fig. 3A and A′). The maximum normalized leak intensities overlapped between the cell lines (SFig. 1D), indicating that leaks in ZO-2 KO and ZO-1 OE cells were of similar magnitude but lasted longer.

**Figure 3.**
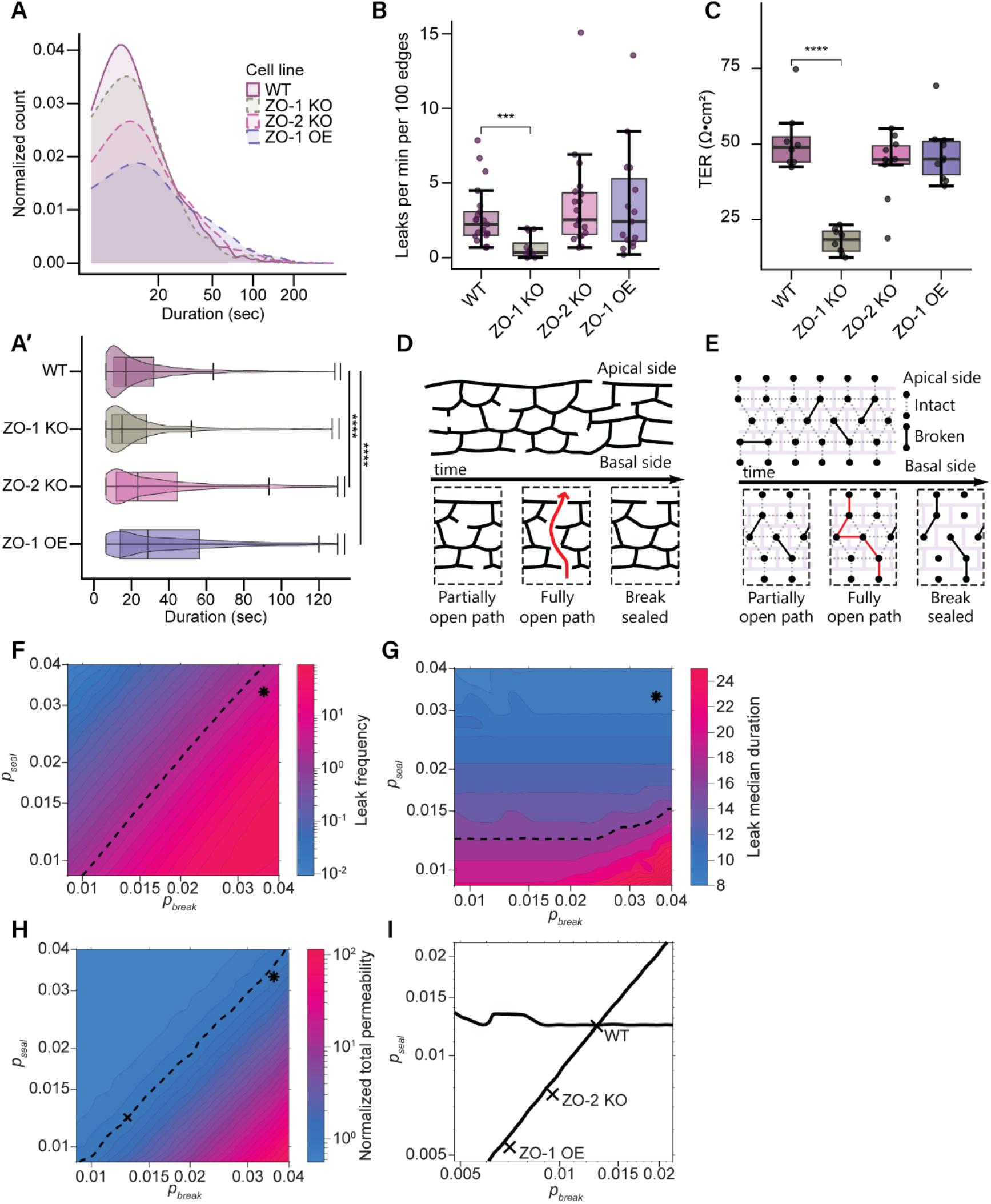
Leak duration, leak frequency and TER, and dynamic strand network simulations. (A) Distribution of leak durations with logarithmic time axis in WT, ZO-1 KO, ZO-2 KO and ZO-1 OE cells. (A′) The same data shown as violin and box plots, with the time axis truncated at 130 s. Leaks lasted longer in ZO-2 KO and ZO-1 OE cells than in WT cells. Leaks still ongoing at the end of the recording were excluded from the duration analysis. (B) Leak frequency, expressed as the number of leaks per 100 bicellular junctions per minute. Each dot represents one video. Fewer leaks were detected in ZO-1 KO cells. (C) TER of the cell lines. Each dot represents one insert. ZO-1 KO cells had lower TER than WT cells. (D) Schematic of the dynamic strand model. Tight junction strands break and seal stochastically, and when breaks in successive strands coincide, a fully open path (red arrow) forms transiently through the network until one of the breaks seals. (E) Corresponding representation in the leak model, in which strand segments between network nodes are either intact or broken. Simulated (F) leak frequency (leaks per 100 junctions per minute) and (G) median leak duration (s) as functions of the break formation rate (p_break_, µm^−1^ s^−1^) and the sealing rate (p_seal_, s^−1^). Dashed lines show the isolines of the experimental WT values. (H) Simulated global permeability of a 547-Da tracer calculated with the TJ model, normalized to the value at the fitted WT rates (cross). Asterisks in F–H mark the rates used in the original TJ model (Tervonen et al., 2019). (I) Intersection of the WT frequency and duration isolines from F and G, giving the fitted WT rates, together with the rates fitted for ZO-2 KO and ZO-1 OE cells. n = 93-2446 leaks (A, A′) and 12-19 videos (B) per cell line, from 4-6 inserts. n = 8-10 inserts per cell line (C). Leak durations and frequencies were compared using Wilcoxon Rank Sum test. TER was compared using t-test. ***p < 0.001, ****p < 0.0001; comparisons without a mark were not significant. Simulated values in F–H are averages over 104 simulations of 2 h for each rate pair.

Leak frequency did not differ between WT (mean 2.71 leaks per 100 junctions per min), ZO-2 KO and ZO-1 OE cells, whereas fewer leaks were detected in ZO-1 KO cells (0.66; p < 0.001; Fig. 3B). Because the leak threshold is relative to the junctional signal, which was highest in ZO-1 KO cells (SFig. 1B), this reduction may partly reflect reduced detection of small leaks rather than fewer leaks.

To relate these junction-level results to global barrier properties, we measured TER. ZO-1 KO cells had markedly lower TER than WT cells (51.4 vs 18.0 Ω·cm^2^; p < 0.0001; Fig. 3C), consistent with their elevated junctional ZnUMBA signal (SFig. 1B). In contrast, TER of ZO-2 KO and ZO-1 OE cells did not differ from WT (Fig. 3C), despite their longer leaks and higher junctional signal.

### Leak frequency and duration constrain strand-level break formation and sealing rates

We next asked whether the observed leaks could arise as transient, fully open paths through a dynamic strand network. In our earlier computational implementation of the dynamic strand model (TJ model), strands break and seal stochastically with a break formation rate (*p*_*break*_) and a sealing rate (*p*_*seal*_), and such open paths occur spontaneously (Fig. 3D) (Tervonen et al., 2019). Because we were interested only in these open paths rather than overall permeability, we simplified the TJ model into a leak model that records the number and duration of fully open paths through the strand network with the given formation and sealing rates (Fig. 3E).

We first simulated the leak model with the mean rates previously fitted to global permeability data from MDCK II cells (*p*_*break*_ = 0.036 µm^−1^ s^−1^ and *p*_*seal*_ = 0.033 s^−1^) (Tervonen et al., 2019), using the average bicellular junction length from our data (7.2 µm) and the same 5 s minimum leak duration as in the experiments. This gave a mean leak frequency of 5.57 leaks per 100 junctions per minute and a median duration of 9.00 s, the same order of magnitude as the experimental WT values (2.71 leaks per 100 junctions per minute and 17.0 s), even though the rates had been derived from global data only.

Global permeability alone cannot determine *p*_*break*_ and *p*_*seal*_ independently, because many rate pairs produce the same permeability. In the original study, *p*_*seal*_ was therefore fixed by assuming a mean strand break duration of 30 s (Tervonen et al., 2019). Leak frequency and duration, however, depend on the two rates differently. In a sensitivity analysis (Fig. 3F), leak frequency increased with increasing *p*_*break*_ and decreasing *p*_*seal*_, was highly sensitive to both, and its isolines followed an approximately constant *p*_*break*_/*p*_*seal*_ ratio. Median duration, in contrast, depended mainly on *p*_*seal*_ (Fig. 3G). Only when *p*_*break*_/*p*_*seal*_ exceeded about 3 µm^−1^ did leak frequency start to decrease and duration increase steeply (Fig. 3F), most likely because frequent, long-lived breaks formed complex multipath openings.

The intersection of the isolines for the experimental WT frequency and median duration gave *p*_*break*_ = 0.013 µm^−1^ s^−1^ and *p*_*seal*_ = 0.012 s^−1^ (Fig. 3I). Both rates are about three times lower than the original values, but their ratio is similar, so the two pairs lie close to the same frequency isoline. This explains why the original values already gave a leak frequency of the right order. The new *p*_*seal*_ corresponds to a mean strand break lifetime of 83.3 s. This is longer than the median leak duration because a fully open path requires several simultaneous breaks and closes as soon as any one of them seals. Assuming an unchanged strand network, fitting the ZO-2 KO and ZO-1 OE data gave lower rates than in WT (ZO-2 KO: *p*_*break*_ = 0.0095 µm^−1^ s^−1^, *p*_*seal*_ = 0.0076 s^−1^; ZO-1 OE: 0.0071 µm^−1^ s^−1^ and 0.0053 s^−1^), shifting them along the WT frequency isoline (Fig. 3I). Thus, the model predicts that in both lines the longer leaks resulted from fewer but longer-lasting strand breaks, with a larger shift in ZO-1 OE. We did not fit the ZO-1 KO data for two reasons: ZO-1 KO has been shown to reduce the number of horizontal strands in MDCK cells (Haas et al., 2022), which violates the model’s assumption of an unchanged network, and its high junctional ZnUMBA signal likely prevented the detection of some leaks (SFig. 1B).

To test how these rate changes affect global barrier properties, we simulated the original TJ model, which includes both bicellular strand dynamics and tricellular tubes, over the same parameter range (Fig. 3H). The permeability of a 547-Da test molecule (polyethylene glycol) with the new WT rates was within 6% of that with the original rates, so the new pair reproduces the global permeability data equally well. Like leak frequency, permeability depended mainly on *p*_*break*_/*p*_*seal*_ and plateaued below a ratio of about 1 µm^−1^, where tricellular tubes became the dominant pathway (SFig. 2A). TER was relatively insensitive to strand dynamics and varied by more than 25% only when *p*_*break*_/*p*_*seal*_ exceeded 2 µm^−1^ (SFig. 2B). The fitted ZO-2 KO and ZO-1 OE rates therefore predict only small changes in TER, consistent with the unchanged TER measured in these lines (Fig. 3C).

In summary, the simulations show that the observed leak frequencies and durations are consistent with stochastic breaking and sealing of tight junction strands, and that a single pair of strand-level rates explains both junction-level leaks and global permeability. Whereas leak frequency, like global permeability, mainly reflects the ratio of the two rates, leak duration provides independent information on the sealing rate that global measurements cannot resolve.

### Leak dynamics are insensitive to myosin II modulation but altered by actin depolymerization

Because ZO-1 links tight junctions to the actin cytoskeleton and leaks in *Xenopus* embryos are repaired by actomyosin contraction (Stephenson et al., 2019), we next asked whether actomyosin activity affects leaks in MDCK II cells. In WT and ZO-1 KO cells, we inhibited myosin II activity with the ROCK inhibitor Y-27632 or the MLCK inhibitor ML-7, promoted cortical myosin II accumulation with 4-hydroxyacetophenone (4-HAP), and depolymerized actin with Latrunculin A (Lat A).

In WT cells, myosin II modulation had little effect on leaks. Leak duration was unchanged by Y-27632 and 4-HAP compared with untreated and DMSO-treated cells, respectively, and ML-7 shortened the median duration by only 2.12 s compared with DMSO (Fig. 4A). None of the myosin II drugs affected leak frequency (Fig. 4B). Lat A did not change leak duration but reduced leak frequency (Fig. 4A and B) and increased the junctional ZnUMBA signal (SFig. 1E). As in ZO-1 KO cells, the lower frequency may partly reflect reduced detection of small leaks against the higher junctional signal. In ZO-1 KO cells, none of the treatments significantly changed leak duration or frequency (Fig. 4D and E).

**Figure 4.**
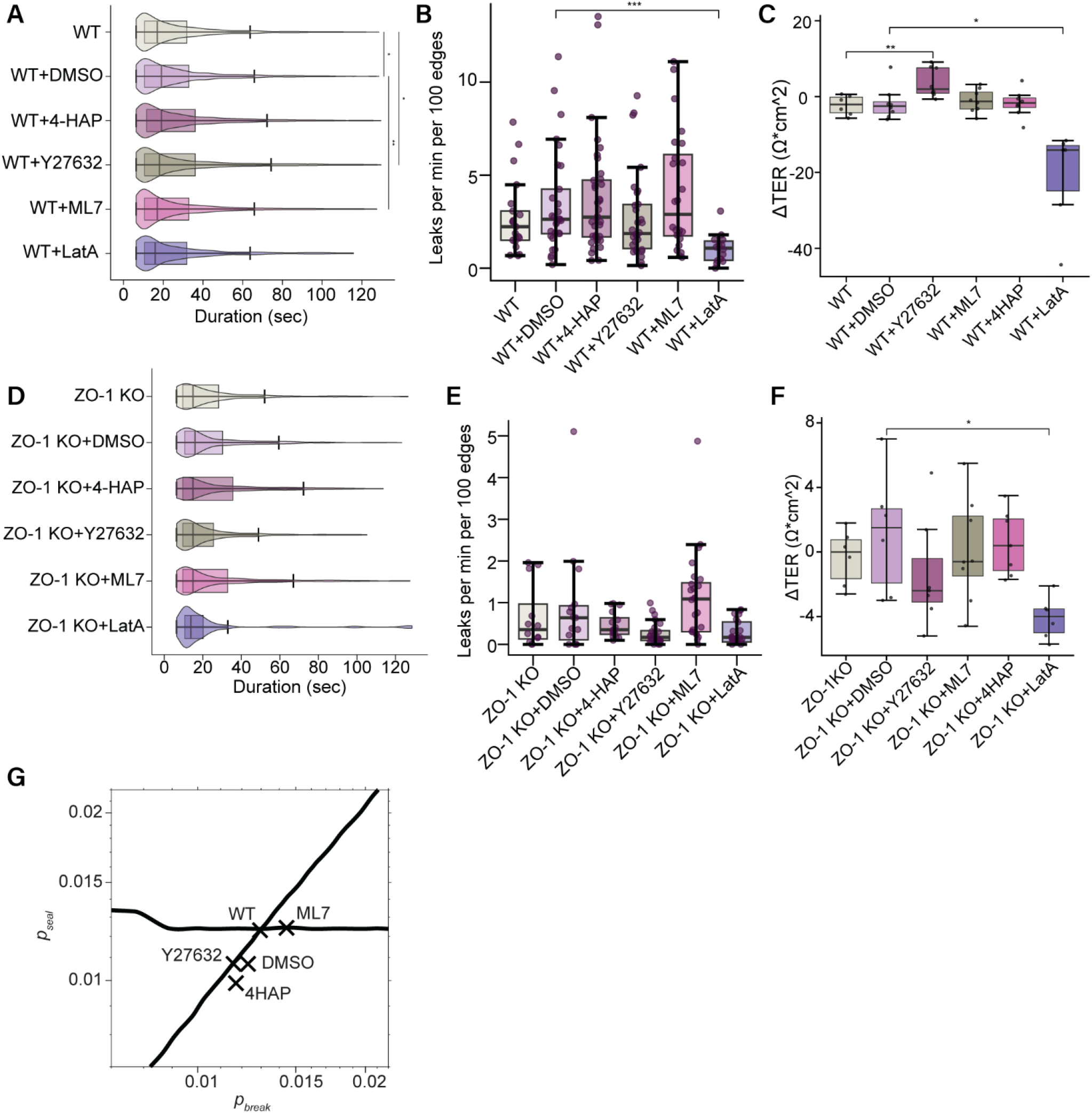
Effects of actomyosin-modulating drugs on leaks and TER in WT and ZO-1 KO cells. WT (A–C) and ZO-1 KO (D–F) cells were treated with the ROCK inhibitor Y-27632 (10 µM), the MLCK inhibitor ML-7 (10 µM), the myosin II activator 4-HAP (5 µM) or Latrunculin A (Lat A; 0.5 µM), or with DMSO as vehicle control. Y-27632 was compared with untreated cells, and the other drugs with DMSO-treated cells. (A and D) Leak durations shown as violin and box plots. Myosin II modulation had only minor effects on leak duration in WT cells (A) and no significant effects in ZO-1 KO cells (D). (B and E) Leak frequency, expressed as the number of leaks per 100 bicellular junctions per minute (a dot represents one video). In WT cells, only Lat A reduced the number of detected leaks (B). None of the treatments changed leak frequency in ZO-1 KO cells (E). (C and F) Change in TER after drug addition (ΔTER = TER after − TER before treatment; a dot represents one insert). In WT cells, Y-27632 increased and Lat A decreased TER (C). In ZO-1 KO cells, only Lat A decreased TER (F). (G) Break formation (p_break_) and sealing (p_seal_) rates fitted for DMSO-, Y-27632-, ML-7-, and 4-HAP-treated WT cells, shown with the WT frequency and duration isolines from Fig. 3I. Lat A was not fitted (see text). n = 142 - 4141 leaks (A, D) and 14 - 36 videos (B, E) per condition, from 4 - 8 inserts. n = 6 - 8 inserts per condition (C, F). Leak durations and frequencies were compared using Wilcoxon Rank Sum test. TER was compared using t-test. * p < 0.05, ** p < 0.01, *** p < 0.001, **** p < 0.0001; comparisons without a mark were not significant.

TER showed that two of the treatments did affect global barrier properties. Y-27632 increased TER in WT cells, as previously reported for MDCK II cells (Van Itallie et al., 2009), but not in ZO-1 KO cells, whereas ML-7 and 4-HAP did not change TER in either line (Fig. 4C and F). Lat A decreased TER in both lines (Fig. 4C and F), consistent with the disruption of tight junctions by actin depolymerization (Shen and Turner, 2005). Thus, Y-27632 changed TER, which mainly reflects ion flux through the pore pathway, without changing leak duration or frequency, indicating that the two can be regulated separately.

Finally, we fitted the leak model to the WT drug data, assuming an unchanged strand network. DMSO and the myosin II drugs shifted the fitted rates only slightly compared with ZO-2 KO and ZO-1 OE, and all remained close to the WT frequency isoline (Fig. 4G). We did not fit the Lat A data, for the same reasons as for ZO-1 KO: actin depolymerization disrupts tight junction structure (Shen and Turner, 2005) and the higher junctional signal likely prevented the detection of some leaks. Together, these results suggest that myosin II activity has little effect on leak dynamics in MDCK II cells, whereas an intact actin cytoskeleton is required to maintain the barrier.

## Discussion

In this study, we used ZnUMBA to detect transient barrier openings, or leaks, in MDCK II cells and to examine how ZO proteins and the actomyosin cytoskeleton affect their number and duration. These leaks were present in MDCK WT cells as well as ZO-1 KO, ZO-2 KO, and ZO-1 OE cell lines. We saw that ZO-1 KO had lower observed leak frequency, higher junctional ZnUMBA signal, and lower TER compared to WT. Furthermore, the observed leaks were longer in ZO-2 KO and ZO-1 OE cell lines than in WT, while TER remained unchanged. We used computational modeling to see if these observed leaks could be explained by the stochastic open paths in the dynamic strand model. The simulations showed that the observed leak frequencies and durations can be explained by stochastic strand breaking and sealing, and that the longer leaks in ZO-2 KO and ZO-1 OE cells correspond to fewer but longer-lasting strand breaks, with only small predicted changes in global permeability and TER. We finally used myosin II modulating drugs that had little effect on the leaks in WT or ZO-1 KO cells. However, actin depolymerization by Lat A led to a decrease in TER in both ZO-1 KO and WT and reduced detected leak frequency in WT.

The leaks observed by Stephenson et al. (2019) in *Xenopus* embryos were initiated by junction elongation and mechanical tension and repaired by Rho flares followed by myosin-driven contraction. Because we did not observe obvious junction elongation at leaking junctions, and myosin II modulation had little effect on leak characteristics, the leaks we observed in MDCK cells likely differ from the tension-induced breaches described in *Xenopus*. Leak duration also differs, further suggesting a distinct phenomenon. In *Xenopus*, leaks last on the order of minutes, whereas our leaks were about an order of magnitude shorter, with a WT median of 17 s, a difference too large to be explained by their longer frame intervals alone. In contrast, the median leak duration in the MDCK ZnUMBA study by Higashi et al. (2022), approximately 30 s, was closer to ours. The remaining difference likely reflects their longer frame interval (20 s vs. our 1.1 s; personal communication), at which most of our leaks would appear in at most one frame, if detected at all. Our simulations showed that stochastic breaking and sealing of tight junction strands, as proposed by the dynamic strand model, is sufficient to explain the leaks we observed. A single pair of strand break formation and sealing rates reproduced both the junction-level leak frequency and duration in WT cells and the global permeability previously measured in MDCK II cells (Tervonen et al., 2019). The model does not specify how breaks are sealed, but sealing appears independent of myosin II and could involve mechanisms such as EpCAM-mediated repair (Higashi et al., 2022).

Our simulations indicate that both the local junctional leak frequency and global permeability depend on the ratio *p*_*break*_/*p*_*seal*_, whereas leak duration depends mostly on *p*_*seal*_. These dependencies break down when breaks are frequent and long-lived (high *p*_*break*_/*p*_*seal*_), as the open paths become complex and contain multiple interconnected routes through the network. Further, the same global permeability can arise from different combinations of leak frequency and duration: either more short leaks or fewer longer leaks. Our results suggest that the longer leaks in ZO-2 KO and ZO-1 OE cells, which occurred at an unchanged frequency, can be explained by fewer but longer-lasting strand-level breaks. Because ZO-1 and ZO-2 nucleate claudin polymerization (Umeda et al., 2006) through concentration-dependent phase separation at the membrane (Beutel et al., 2019; Sun et al., 2025), their levels could plausibly affect how often strands break and how quickly they seal. The simulations further predict that these changes should not affect global permeability or TER, and our TER measurements confirmed the latter. Consistent with this, earlier studies found no obvious barrier defects after ZO-2 KO (Belardi et al., 2020; Otani et al., 2019) or ZO-2 KD (Van Itallie et al., 2009) in MDCK cells, and expressing ZO-1 above endogenous levels did not change global permeability (Van Itallie et al., 2009). Tokuda et al. (2014) reported increased TER and 4-kDa dextran flux in one ZO-1 KO rescue clone that expressed ZO-1 well above endogenous levels, but their overexpression was likely considerably higher than ours (1.23-fold) or that of Van Itallie et al. (2009) (1.35-fold), and they used a larger tracer, so their results are not directly comparable.

Loss of ZO-1 changed the leaks in our cells. Leak duration was similar to WT, but fewer leaks were detected, and the junctional ZnUMBA signal was markedly higher. ZO-1 KO in MDCK cells reduces the number of horizontal strands, i.e. the strands a molecule has to cross, from approximately four to two (Haas et al., 2022). In the dynamic strand model, fewer strands make open paths through the network much more likely. When such paths become frequent and long-lived, they would merge into persistent openings that appear as an elevated junctional signal rather than as discrete leaks, which could explain both the higher signal and the lower leak frequency. In addition, the higher signal likely masked some smaller leaks, because the leak threshold is relative to the baseline.

ZO-1 KO also reduced TER by approximately 65%. Previous ZO-1 KO and KD studies reported increased leak-pathway permeability, measured as the flux of larger PEGs (Van Itallie et al., 2009) or of 4-kDa dextran (Haas et al., 2022; Tokuda et al., 2014), but only small or no changes in TER, or even an increase (Belardi et al., 2020; Haas et al., 2022; Tokuda et al., 2014; Umeda et al., 2006; Van Itallie et al., 2009). Fewer strands acting as resistors in series should lower TER, whereas the reduced junctional claudin-2, seen both in our cells and by Haas et al. (2022), should partly compensate. Why the TER reduction was larger in our cells than in previous studies remains unclear, but differences in culture conditions, such as inverted seeding on 1 µm filters, may contribute.

Because ZO-1 links tight junctions to the actomyosin cytoskeleton via its C-terminus (Itoh et al., 1997; Schwayer et al., 2019; Spadaro et al., 2017; Van Itallie et al., 2017), we expected myosin II modulation to affect leaks, but it had little effect. Y-27632 nevertheless slightly increased TER in WT cells, as previously reported (Van Itallie et al., 2009), without affecting leaks, indicating that TER and leaks can be regulated separately. ML-7 and 4-HAP, in contrast, had no detectable effect on either TER or leaks. MLCK inhibition mainly increases TER when MLCK has been activated (Turner et al., 1997; Yu et al., 2010), which may explain why ML-7 had no effect in our unstimulated cells. However, because we lacked an independent readout of drug activity, we cannot exclude that ML-7 and 4-HAP were ineffective under our conditions. Lat A decreased TER and increased the junctional signal, resembling our ZO-1 KO phenotype. This is consistent with Shen and Turner (2005), who showed that Lat A reduces TER in MDCK cells within minutes through caveolae-mediated endocytosis of occludin, and suggests that an intact actin cytoskeleton is required for the barrier.

The junctional ZnUMBA signal in our cells likely reflects, at least in part, Zn^2+^ permeating the tight junctions through claudin-2 pores, possibly together with stepwise diffusion through the strand network and leaks too small to be resolved. Higashi et al. (2022) performed ZnUMBA in claudin-2 KO MDCK II cells, which lack our background signal and therefore allow lower-intensity leaks to be detected, including leaks that the higher junctional signal may have masked in our data. We show, however, that ZnUMBA can also be applied to MDCK II cells that express claudin-2. Our imaging parameters also differed. Our shorter frame interval allowed us to resolve short leaks, whereas their longer recordings (over 30 min vs. our 10 min) capture long leaks that in our videos would often still be ongoing at the end of the recording and were therefore excluded from the duration analysis. The two approaches are thus complementary, each detecting leaks that the other is likely to miss. Shorter frame intervals, together with shorter filtering windows in the analysis, could reveal even shorter leaks in the future.

Our results concern ions and small solutes. ZnUMBA reports the passage of Zn^2+^, the apical FZ3 signal shows that the ∼0.8-kDa indicator also crosses the barrier, and the model was compared with permeability data for a 547-Da tracer. Our results are therefore most relevant for solutes that are too large to pass through claudin pores but smaller than 1 kDa. Whether larger, multi-kDa molecules use the same transient open paths remains to be determined, and comparisons with previous studies using larger tracers, should be interpreted with caution.

Our leak analysis has some limitations. Because the leak threshold is defined relative to the local baseline, leaks are harder to detect when the junctional signal is high, which likely led to an underestimation of leak frequency in ZO-1 KO and Lat A-treated cells. The measured leak duration also differs from the duration of the underlying open path: it includes the time needed for Zn^2+^-FZ3 to clear from the lateral intercellular space, which was not included in the model. Finally, leaks still ongoing at the end of the 10-min recordings were excluded from the leak duration analysis, which biases durations downward. Our model also assumes a uniform strand network, although real networks are not uniform: their complexity depends on tight junction protein composition (Saito et al., 2023), and strands in MDCK II cells are often discontinuous (Furuse et al., 2001). The model therefore represents an average network. We also assumed that ZO manipulations and drug treatments change only the break formation and sealing rates, not the network structure, which is why ZO-1 KO and Lat A were not fitted. Nevertheless, the good agreement between the models, our leak data and previous global permeability data suggests that the model captures the key mechanisms of the leak pathway.

## Conclusion

In conclusion, fast ZnUMBA imaging revealed frequent, short-lived leaks for ions and small solutes at bicellular junctions of claudin-2-expressing MDCK II cells. Their frequency and duration were consistent with open paths formed by the stochastic breaking and sealing of tight junction strands rather than myosin-driven repair, and the same strand-level rates explained both the junction-level leaks and global permeability. Altered ZO protein levels changed these strand dynamics. ZO-2 KO and ZO-1 OE prolonged leaks without affecting leak frequency or TER, a change that global barrier measurements alone would not detect. Our approach combining ZnUMBA with computational modelling therefore offers a way to link tight junction strand dynamics to epithelial barrier function.

## Methods

### Cell culture

MDCK II ZO-1 KO and MDCK ZO-2 KO cell lines were kindly sent by Briand Belardi and Daniel A. Fletcher (University of California, Berkeley, USA). MDCK II WT cells, MDCK II cells expressing ZO-1 fused to mRuby fluorescent protein, MDCK II ZO-1 KO cells, MDCK II ZO-1 KO cells transiently transfected with mCherry-actin-chromobody, and MDCK II ZO-2 KO cells, and MDCK ZO-2 KO cells transiently transfected with mCherry-actin-chromobody were maintained in regular cell culture conditions, using MEM (Gibco) with Earle’s salts and GlutaMAX, and supplemented with 10 % fetal bovine serum (FBS), and 1% penicillin/streptomycin (ZO-1 KO cell lines) or 0.25 mg/ml G-418 (Roche) (ZO-1 mRuby, ZO-1 KO actin-chromobody-mCherry, and ZO-2 KO actin-chromobody-mCherry), and kept in 5 % CO_2_ concentration at +37 °C. 0.25 mg/ml G-418 (Roche) Cells were subcultured every 1 to 2 weeks and the medium was changed every 5^th^ day. WT MDCK ZO-1 mRuby cells were chosen to visualize the location of the tight junctions. The motive to use actin chromobody on KO cell lines was to better visualize the cells, but this turned out not to be necessary. ZnUMBA data shown in Fig. 2, 3, and 4 denoted as ZO-1 KO or ZO-2 KO used the ZO-1 KO actin-chromobody-mCherry line or ZO-2 KO actin-chromobody-mCherry line, respectively. Immunostaining images, TER measurements and drug treatments were performed using ZO-1 KO and/or ZO-2 KO cell lines without actin-chromobody-mCherry expression.

Following Higashi et al. (2023), samples for ZnUMBA were prepared using inverted seeding. The outside surface of transparent, 24-well inserts with 1.0 µm pore size (Sarstedt) was coated by incubating 0.1 mg/ml collagen-I solution for approximately 45 min at room temperature, of which 30 min were under ultraviolet light. Cells were detached from a 25 cm^2^ flask 0.25 % trypsin with EDTA and incubated at +37 °C for 15-20 min. Approximately 1.4×10^5^-2.2×10^5^ cells were seeded on the inverted insert and incubated at +37 °C for 50-60 min. After the incubation, the inserts were placed to 24-wells and cultured for 5 days, with a medium change on day 3.

### ZnUMBA imaging

ZnUMBA imaging was performed using imaging medium (1xMEM (Gibco), 1% penicillin-streptomycin, 5% FBS, without phenyl red) with either 2 mM ZnCl_2_ (Sigma-Aldrich) or 10 µM FluoZin-3 indicator (FZ3, cell impermeable, Invitrogen) added to the apical and basal sides, respectively. For imaging, Nikon A1R with 40x water immersion objective (lambda S, apochromat, DIC N2, numerical aperture 1.15, working distance 0.61 mm, Nikon) was used. The stage top incubator was set to +37 °C and 5% CO_2_ gas flow was used during imaging to prevent pH and temperature changes. FZ3 was excited using 488 nm laser (detector HV: 70, laser intensity: 0.30 %), 10-minute time-lapses were taken using galvano scanning with pixel dwell time of 2.2 µs and frame interval of 1.06 s (0.940 FPS). Pinhole size was 30.7µm, image resolution was 512x512, pixel size was 0.25 µm and 2.5 zoom was used. Perfect focus, a feature in A1R that helps to keep the focus constant, was used. From each sample, 3-5 videos were taken. ZO-1 mRuby was excited using 561 nm laser (detector HV: 90, laser intensity: 0.5 %) and captured only for tight junction visualization purposes. For drug treatment, WT or ZO-1 KO cells were incubated with 10 µM Y-27632 (Stemcell Technologies), 10 µM ML-7 (Bio-Techne Sales Corp.), 5 µM 4-HAP (Sigma-Alrich) or 0,05% DMSO for 1 h, or 0.5 µM LatA (Tocris) for 30 min in + 37 °C before imaging and maintaining the drug treatment during imaging.

### Immunostainings, imaging and flow cytometry

MDCK cells grown on inserts were fixed either with 4% paraformaldehyde (PFA) for 10 min, or methanol for 5 min, and washed 2x with PBS. Then, PFA-fixed samples were permeabilized for 10 min in PBS with 0.1% Triton X and 0.5% BSA. Primary antibodies for PFA-fixed samples (1:200 ZO-1 rabbit (31-7300), 1:200 claudin-2 mouse (32-5600)) were diluted in 3 % BSA and incubated overnight in +4 °C. For methanol-fixed samples, permeabilization was not performed and primary antibodies (1:200 ZO-1 mouse (33-9100), 1:100 ZO-2 rabbit (71-1400)) were diluted in 3 % BSA and incubated in +4 °C overnight. After primary antibody incubation, permeabilization was repeated for samples with both fixation methods, and secondary antibodies (For PFA fixed samples: 1:200 AlexaFluor 488 anti-mouse (A11029), 1:200 AlexaFluor 568 anti-rabbit (A11011), and 1:100 ATTO phalloidin 643, for methanol fixed samples: 1:200 AlexaFluor 488 anti-rabbit (A11008) and 1:200 AlexaFluor 568 anti-mouse (A11031)), diluted in 3% BSA were incubated 1h in room temperature protected from light and washed once with water and once with PBS. All antibodies from Invitrogen and phalloidin from Atto-Tec. Samples were mounted using Prolong Diamond Antifade with DAPI and imaged using Evident FV4000 confocal microscope using 60x oil objective (Olympus UPlanXApo 60x/1.42, WD 0.15 mm).

Flow cytometry analysis was done by fixing suspended WT and ZO-1 mRuby cell with 4% PFA for 10 mins. Cells were washed once before 10 min permeabilization with 0.5% Triton X and 0.5% BSA, after with primary antibodies (1:200 ZO-1 mouse (33-9100) Invitrogen, 1:200 vimentin rabbit (ab92547), Abcam) were incubated 30min in room temperature. Next, cells were washed once with PBS before 30 min incubation with the secondary antibodies (1:200 AlexaFluor 488 anti-mouse (A11029), 1:200 AlexaFluor 647 anti-rabbit (A21245), Invitrogen). After immunostaining, cells were suspended in PBS with 1 mM EDTA for flow cytometry analysis. Using Beckman Coulter CytoFlex S flow cytometry, ZO-1 mRuby intensity and immunolabeled ZO-1 intensity were measured from WT and ZO-1 mRuby cells and median immunolabeled ZO-1 fluorescence intensity was compared between the cell lines. Overexpression level was counted as ZO-1 mRuby positive cells’ labeled ZO-1 median intensity divided by WT cells’ labeled ZO-1 median intensity.

### TER measurements

TER was measured using Millicell ERS-2 Voltohmmeter in RT. Inserts with cells cultured for 5 days were measured. One insert was measured three times with approximately 10 min interwall between measurement rounds. The final TER values reported was the average of the three measurements. Between measurement rounds cells and electrode were kept at +37 °C. When measuring effects of DMSO (0.05 %), Y-27632 (10 µM), 4-HAP (5 µM), ML-7 (10 µM) or Lat A (0.5 µM) treatment, TER was measured before and after the drug treatment. Incubation at +37 °C was 1h for other drugs except Lat A, for which it was 30 min.

### Image analysis and leak detection

Tissue barrier function monitored with ZnUMBA was captured as 2D time-lapse videos. Before analysis, the videos were aligned using the StackReg plug-in in ImageJ (version 1.54f) (Thevenaz et al., 1998). To account for cell drift, cells were segmented using Cellpose (Stringer et al., 2021) every 10 frames and tracked throughout the videos. Subsequent analyses were conducted using a custom program in MATLAB (R2023a, update 2; MathWorks Inc.). Each pair of connected cells (with connected cell masks) yields a unique cell edge, the mask of which was determined by dilating the two cells’ common boundary to a width of 7 pixels (∼3 µm). Edges shorter than 10 pixels were excluded. Edge intensities were extracted as the average ZnUMBA intensity within the respective edge mask in each frame.

To avoid detecting leaks from random signal fluctuations and measurement noise, we first applied a 10-s moving-average filter to the edge intensity traces. To correct for drifts in the whole-tissue ZnUMBA signal over the imaging period, we defined a baseline at each time point as the lowest intensity within a 50-s window around that time point. The leaks were defined as continuous periods in which the ZnUMBA signal was at least 20% above this baseline for at least 5 s. Successive leaks separated by a single frame were joined to account for sporadic noise. For each leak, we extracted its intensity relative to the baseline and its duration. For comparison between cell lines, intensity values were normalized to the mean intensity of the edge excluding values above the leak threshold.

To test whether leaks involved the whole junction or only part of it, we divided each edge mask into five equal subsections along its length. If, during a leak, the summed intensity in the 1st and 2nd subsections differed by at least 10% from that in the 4th and 5th subsections, the leak was classified as localized to the corresponding end of the junction. Otherwise, it was classified as a whole-junction leak.

### Computational modeling

Two computational models were used in the study. The TJ model presented by Tervonen et al. (2019), which describes the effect of strand-level dynamics on the global epithelial tracer permeability and TER (https://github.com/atervn/tj_model). For a detailed description of the model see Tervonen et al. (2019). Briefly, the model describes the tight junction strand network as a collection of compartments with transient boundaries describing the strands. Breaks are formed and sealed during the simulation given break formation and sealing rates, respectively. This model is referred as the *TJ model*.

We developed another, more simplified model based on the original TJ model to focus solely on leaks through the tight junction strand network rather than the permeability values. This model is referred as the *leak model*. The model is described using a graph, with vertices representing the compartments lined by the strands and edges representing the connections between the compartments. The edges have two states: active or inactive, describing the broken or intact states of the strand between two compartments. Two additional rows of vertices were added on the basal and apical sides of the strand network to describe the respective compartments and connected only to the closest vertex within the network. During the simulation, the state of the edges is changed based on the break formation and sealing rates, as in the TJ model. All transient fully open path between sources and sinks were detected using the Breadth-First search algorithm, and the found leaks, including their quantity, start, and end times, were recorded. The model was simulated 10^4^ times for each set of break formation and sealing rate pairs to achieve reliable average results. The model is presented in detail in SText 1 and the code is available in Zenodo (Tervonen, 2026) and Github (https://github.com/atervn/leak_model). Both models were implemented with MATLAB.

### Statistical analysis and plotting

Rstudio (version 2023.12.1) with packages tidyverse (version 2.0.0), readxl (version 1.4.3), ggthemes (version 5.1.0), R.matlab (version 3.7.0), svglite (version 2.1.3), ggsignif (version 0.6.4), plyr (1.8.9) and ggplot2 (version 3.5.0) were used for statistical analysis and for plotting for the experimental data. Because data was not normally distributed, Kruskal-Wallis rank sum test was used to test for differences between multiple variables, and pairwise Wilcoxon rank sum test with Holm–Bonferroni p-value adjustment was used to test differences between two samples. T-test with Holm–Bonferroni p-value adjustment was used to compare two sample groups in TER data. To produce plots, ggplot2-package was used. Simulation results were plotted using MATLAB.

## Acknowledgements

We acknowledge the Tampere Imaging Facility (TIF) and Flow Cytometry Facility for services, and Riikka Järvinen for her help with the FACS experiments. We would also like to acknowledge Brian Belardi and Daniel A. Fletcher for providing the ZO-1 KO and ZO-2 KO cell lines.

## Author contributions

Data curation: A.P., A.T.; Formal analysis: A.P., A.T.; Funding acquisition: T.O.I., S.N., H.T.; Investigation: A.P., A.T.; Methodology: A.P., A.T., H.T.; Project administration: T.O.I., S.N., A.T.; Resources: T.O.I., S.N., H.T.; Visualization: A.P., A.T.; Writing – original draft: A.P., A.T., H.T.; Writing – review & editing: A.P., A.T., H.T., S.N., T.O.I.

